# Sensory cilia act as a specialized venue for regulated EV biogenesis and signaling

**DOI:** 10.1101/2021.02.04.429799

**Authors:** Juan Wang, Inna A. Nikonorova, Malan Silva, Jonathon D. Walsh, Peter Tilton, Amanda Gu, Maureen M. Barr

## Abstract

Extracellular vesicles play major roles in intercellular signaling, yet fundamental aspects of their biology remain poorly understood. Ciliary EV shedding is evolutionary conserved. Here we use super resolution, real time imaging of fluorescent-protein tagged EV cargo combined with *in vivo* bioassays to study signaling EVs in *C. elegans*. We find that neuronal sensory cilia shed the TRP polycystin-2 channel PKD-2::GFP-carrying EVs from two distinct sites - the ciliary tip and the ciliary base. Ciliary tip shedding requires distal ciliary enrichment of PKD-2 by the myristoylated coiled-coil protein CIL-7. Kinesin-3 KLP-6 and intraflagellar transport (IFT) kinesin-2 motors are also required for ciliary tip EV shedding. Blocking ciliary tip shedding results in excessive EV shedding from the base. Finally, we demonstrate that *C. elegans* male ciliated neurons modulate EV cargo composition in response to sensory stimulation by hermaphrodite mating partners. Overall, our study indicates that the cilium and its trafficking machinery act as a specialized venue for regulated EV biogenesis and signaling.

## Main

Extracellular vesicles (EVs) were once considered cellular debris^1–3^ but are now recognized as a powerful mode of cellular crosstalk in different physiological and pathological states^4,5^. Likewise, the cilium went from being viewed as a vestigial organelle to a critical signaling nexus. Cilia receive extracellular cues and also shed EVs^6–12^. Ciliary EVs provide a cellular and organismal conversation system, providing a unique opportunity to decode the language of EVs. In *Chlamydomonas* and *C. elegans*, ciliary EVs act as signaling devices^13,14^. In cultured mammalian cells, ciliary EVs regulate ciliary disposal but also receptor abundance and signaling, ciliary length, and ciliary membrane dynamics^9–11,15^. Mammalian cilia appear to produce EVs from the tip and along the ciliary membrane^16,17^. However, the functional significance of shedding at the distinct locations, as well as molecular mechanisms that regulate ciliary EV biogenesis remain elusive.

Autosomal dominant polycystic kidney disease gene product, polycystin-2, localizes to cilia and extracellular vesicles (EVs) throughout evolution. Loss of ciliary localization of polycystin-2 leads to cystogenesis in mouse model ^18^. The relationship between polycystin-2 ciliary localization and EV biogenesis represents a knowledge gap in the field. *C. elegans* ciliated sensory neurons release environmental EVs carrying PKD-2::GFP as visualized by live imaging. EVs are also shed at the ciliary base into the lumenal compartment formed by glial cells enwrapping neuron receptive endings as visualized by transmission electron microscopy and tomography ^14,19^. The sites and mechanisms regulating ciliary EV biogenesis are not known.

To study ciliary EV biogenesis and shedding, we used super-resolution microscopy to visualize fluorescently-labeled EV cargo in real time from living animals at 120nm 2D resolution (compared to 200-300nm with conventional imaging). We generated transgenic animals co-expressing PKD-2::GFP with markers that labeled the ciliary axoneme (β-tubulin TBB-4::tdTomato) or transition zone (nephrocystin-1 NPHP-1::dsRed). Analysis of PKD-2::GFP distribution relative to the markers of ciliary compartments revealed two characteristic patterns of the PKD-2 distribution. In the first pattern, environmentally released PKD-2::GFP-carrying EVs were observed outside the male (Fig. 1 a – b, white arrowheads). The presence of environmental EVs was accompanied by PKD-2::GFP enriched at ciliary tip and distributed along the ciliary membrane and periciliary membrane (PCM) at the ciliary base (Figure 1 a - b). In this first pattern, we observed bursts of EVs being shed from the ciliary tip. In the second pattern, ciliary base EVs carrying PKD-2::GFP EVs extended from a protrusion at the PCM below the ciliary transition zone, with no or very little environmental EVs observed (Figure 1 c-d). Ciliary base EVs contained both PKD-2::GFP and TBB-4::tdTomato or NPHP-1::dsRed whereas environmentally released EVs did not carry TBB-4 or NPHP-1. TBB-4::tdTomato and NPHP-1::dsRed fluorescence in ciliary base EVs was continuous with PCM (Fig. 1 c), suggesting that ciliary base EVs were captured in the process of budding from PCM. These data taken together suggested that cilia shed PKD-2 EVs at two sites: the ciliary tip and the PCM at the ciliary base (Fig. 1e).

**Figure 1:**
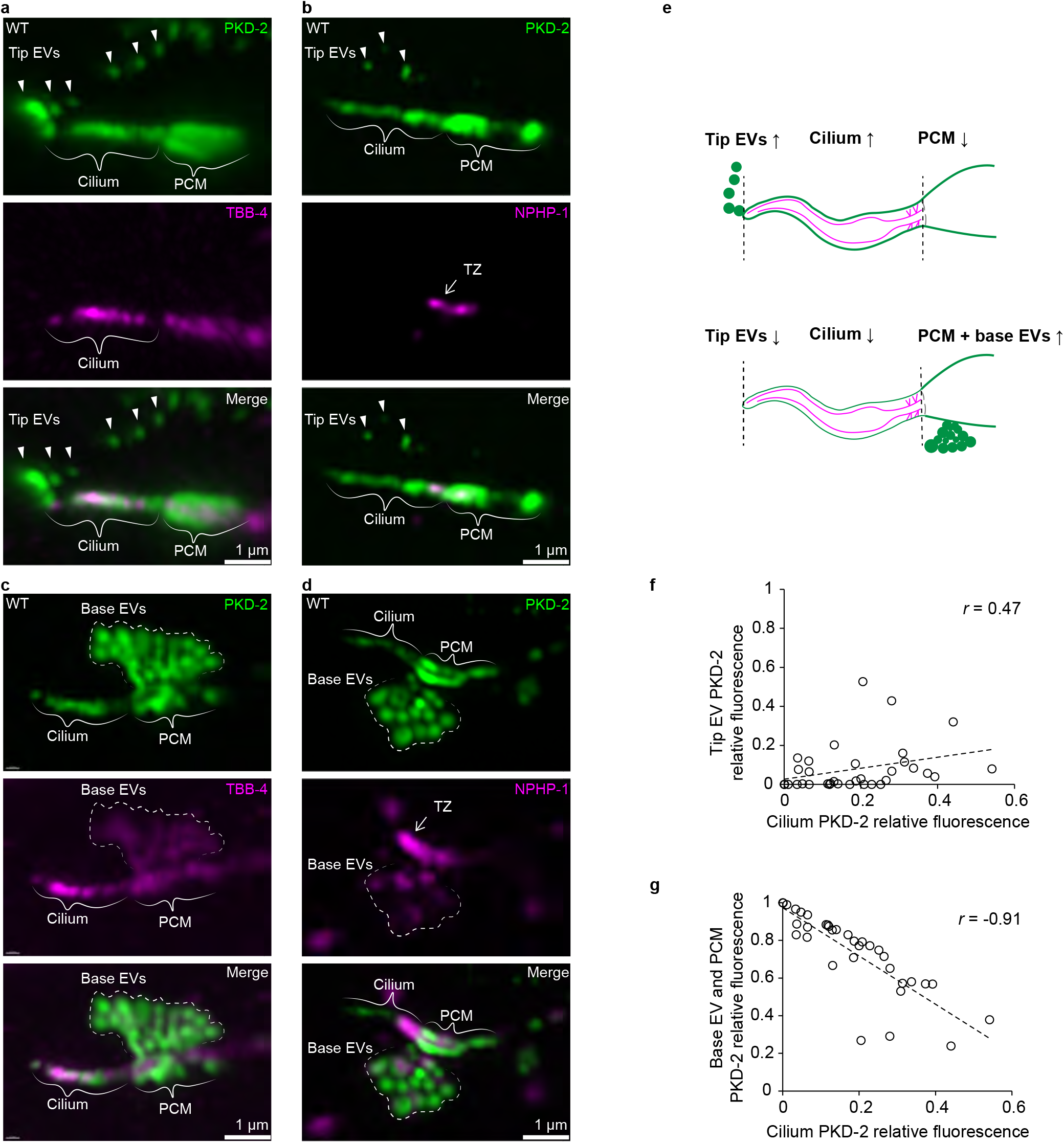
PKD-2 EVs are shed from either ciliary tip or ciliary base of the <u>ce</u>phalic <u>m</u>ale CEM cilium. **a-d** PKD-2:GFP EVs captured at the moment of shedding from the tip (in **a** and **b)** or from the base (in **c** and **d**) of the CEM cilium. Axoneme is labeled with β-tubulin TBB-4::tdTomato, transition zone is labeled with NPHP-1::dsRed. White arrowheads indicate environmental EVs outside the animal, white arrows point to the transition zone. Dashed white lines outline base EVs in the process of shedding, solid white lines indicate PCM, <u>p</u>eri<u>c</u>iliary <u>m</u>embrane this is situated below the transition zone. **e**, Schematic cartoon summarizing antagonistic relationship between PKD-2 EVs shed from the tip and from the base. **f-g**, Correlation plots showing that fluorescence intensity of the tip PKD-2::GFP EVs positively correlates with that of the ciliary shaft (**f**), whereas fluorescent intensity of the base PKD-2::GFP EVs inversely correlates with the presence of PKD-2::GFP at the ciliary shaft (**g**). Spearman test, 35 data pairs. *r* = 0.47, *p* = 0.004 for (**f**), *r* = −0.91, *p* < 0.0001 for (**g**).

To explore the relationship between EVs shed from the ciliary tip and from the ciliary base, we quantified the relative distribution of PKD-2::GFP between the environmental EVs from the tip, the ciliary membrane, and the base. Enrichment of PKD-2::GFP along the ciliary membrane positively correlated with the quantity of the environmental EVs, suggesting that PKD-2::GFP is incorporated into the ciliary membrane in order to be shed in environmental EVs. On the other hand, the relative amount of PKD-2::GFP at the ciliary base inversely correlated with the presence of PKD-2::GFP in the ciliary membrane. These data suggest that environmental EV biogenesis involves a mechanism for translocating PKD-2::GFP from the PCM to the ciliary tip.

To identify mechanisms driving PKD-2 protein to the ciliary tip, we focused on two genes required for environmental release of PKD-2::GFP EVs but not for ciliogenesis: the myristoylated coiled-coil protein CIL-7 and the kinesin-3 KLP-6. In wild-type, PKD-2 distributed along the length of the cilium and was enriched at the ciliary tip (Fig. 2 a). In the *cil-7* mutant, PKD-2::GFP was not delivered to the ciliary tip as evidenced by the absence of PKD-2::GFP at the most distal region of the cilium (Fig. 2 a-b). We also observed an order of magnitude higher accumulation of PKD-2::GFP at the ciliary base relative to the ciliary membrane (Fig. 2 c). In *cil-7* mutant cilia, the length of the PKD-2 distal exclusion zone ranged from 250 nm to 700 nm (Fig. 2 d-f). In the absence of CIL-7, PKD-2 accumulates at the proximal ciliary membrane and ciliary base, suggesting that CIL-7 is required to locate PKD-2 to distal cilium and ciliary tip.

**Figure 2.**
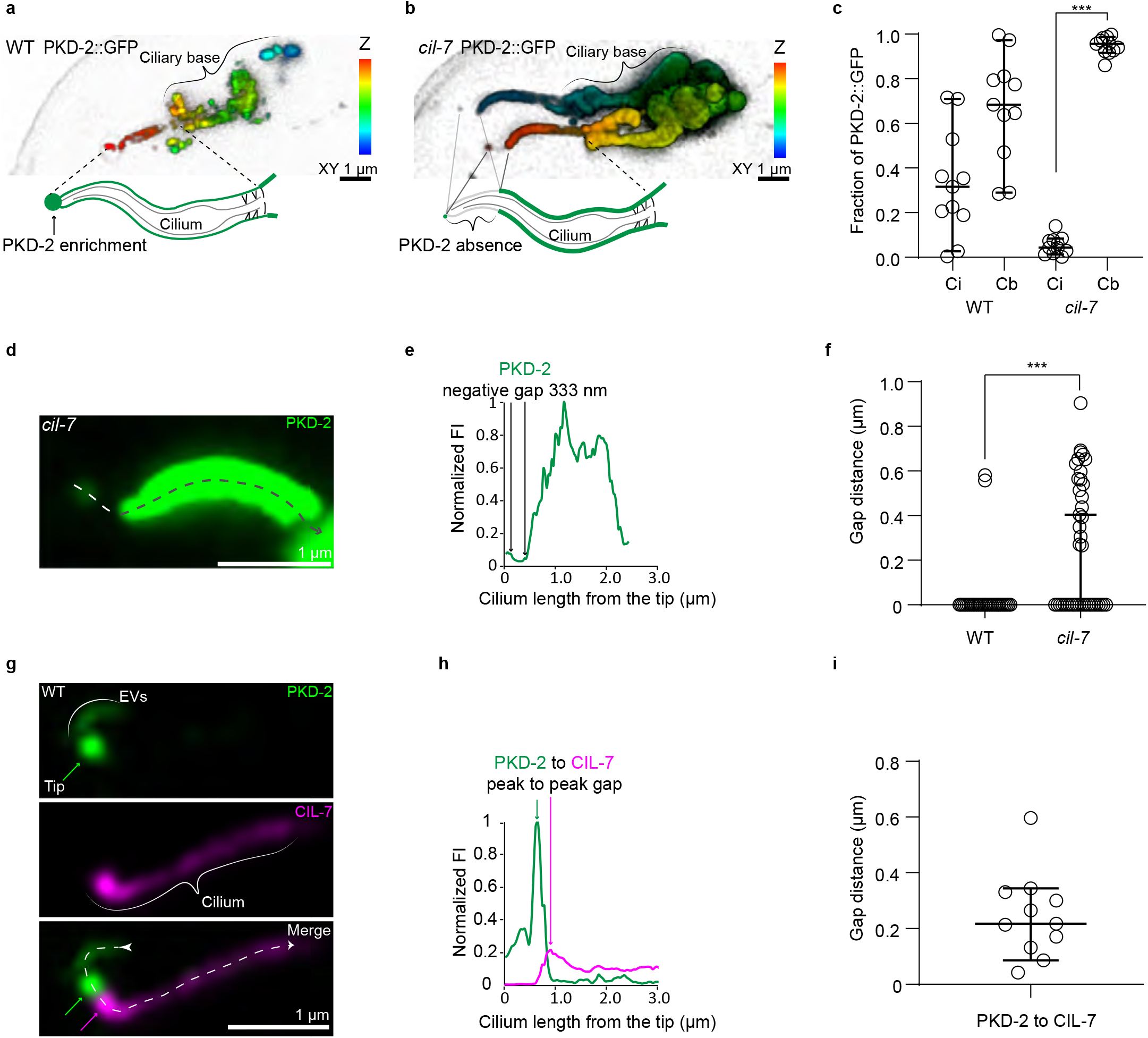
PKD-2 release from the tip of CEM cilium requires its distal ciliary enrichment mediated by CIL-7. **a-b**, Depth coded 3D projections of PKD-2::GFP distribution in the CEM cilia in wild type and the *cil-7* mutant respectively. In the wild type, PKD-2 is distributed evenly along the cilium including ciliary base, shaft, and tip. The *cil-7* mutant cilium lacks PKD-2::GFP at the tip and at the distal region, whereas considerable accumulations of PKD-2::GFP are observed at the ciliary shaft and the ciliary base. **c,** Measured PKD-2::GFP relative fluorescence intensity distribution along ciliary shaft (Ci) and ciliary base of wild type and the *cil-7* mutant cilia. The *cil-7* mutant demonstrates heavily skewed distribution of PKD-2 with significant accumulations at the ciliary base. \*\*\**p*<0.001 by Kruskal-Wallis test with Dunn’s multiple comparison, n=11 wild type and 11 *cil-7* mutant cilia. **d-e**, Close-up image of PKD-2::GFP distribution along the *cil-7* mutant cilium. Dashed line with arrowhead indicates direction of fluorescent intensity profiling depicted in **e**. **f**, The PKD-2 exclusion zone in the *cil-7* mutant reaches up to 1 μm, which is significantly larger than most wild-type cilia. \*\*\**p*<0.001 by Mann-Whitney test, n=31 wild type and 42 *cil-7* mutant cilia. **g,** Representative image of the wild-type CEM cilium showing PKD-2::GFP enrichment at the ciliary tip. Newly formed string of tip EVs is labeled as EVs and indicated by white line; it extends from the PKD-2 enriched ciliary tip to above focal planes. Green and magenta arrows point to the corresponding positions of PKD-2 and CIL-7 peaks shown in **h**. **h**, Fluorescent intensity profiling of the cilium from panel **g** shows that PKD-2 and CIL-7 are enriched at distinct locations positioned at 256 nm apart from each other. **i**, Distances measured between PKD-2 and CIL-7 enrichment areas at the distal parts of wild-type cilia, n=11. All the plots show median values with 95% confidence intervals.

To test whether CIL-7 functioned at the distal ciliary compartment, we imaged the CIL-7::tagRFP reporter co-expressed with PKD-2::GFP (Fig. 2 g). Lengthwise fluorescent profiling along the double-labeled cilia (from the very tip toward the proximal region) revealed that both PKD-2::GFP and CIL-7::tagRFP fluorescent signals peaked exclusively at the area adjacent to the ciliary tip (Fig. 2g). More specifically, PKD-2::GFP peaks were always accompanied by the presence of the CIL-7::tagRFP peaks with the average distance of 246 nm between them (Fig. 2 g-i, Extended data 1 a-b (cilia #1 through #4)). The distance between the PKD-2 and CIL-7 peaks at the ciliary tip corresponded to the length of the PKD-2::GFP exclusion zone in the *cil-7* mutant (Fig. 2 d-f). These data suggest that delivery of PKD-2 to the ciliary membrane is not sufficient for its enrichment at the ciliary tip. Nearly complete absence of PKD-2::GFP at the ciliary tip of the *cil-7* mutant suggested that CIL-7 acted at the distal region of the cilium facilitating enrichment of PKD-2::GFP at the ciliary tip for its exit as environmental EVs.

How does CIL-7 presence at the distal cilium influence enrichment of PKD-2 at the ciliary tip? CIL-7 is a cargo of environmental EVs^14,20^ and may be involved the recruitment of PKD-2 into EVs containing CIL-7. To test this, we determined whether CIL-7 and PKD-2 were co-sorted into EVs and found PKD-2 and CIL-7 mostly on separate EVs (Extended data 1 a-b). Super-resolution imaging revealed three scenarios of PKD-2 and CIL-7 EV shedding: (i) CIL-7 EV budding from the ciliary tip without accompanying PKD-2 (Extended data 1 b), (ii) PKD-2 EV budding from the CIL-7 enriched area of the ciliary tip (as depicted on Fig. 2 c, Extended data 2), and (iii) several PKD-2 EVs budding simultaneously from a large CIL-7 protrusion at the ciliary tip and separation into PKD-2 and CIL-7 carrying EVs (Extended data 3). Formation of the PKD-2 EVs at the ciliary tip was always accompanied by CIL-7 distal ciliary enrichment, whereas budding of CIL-7 EVs was independent of PKD-2 presence. Combined, these data suggest that CIL-7 acts an EV launching pad by enriching PKD-2 at the ciliary tip and packaging PKD-2 into environmental EVs.

The continuous loss of CIL-7 from the ciliary tip during PKD-2 EV shedding suggests that distal ciliary pool of the CIL-7 protein has to be maintained. Thus, we next sought to identify the mechanisms regulating PKD-2 and CIL-7 distribution in cilia. The cell-specific ciliary kinesin-3 KLP-6 is important for PKD-2 localization in cilia and in EVs but not for ciliogenesis^14,21,22^. We compared ciliary localization of PKD-2 and CIL-7 in wild type and *klp-6* mutants at the onset of PKD-2::GFP and CIL-7::tagRFP expression in the fourth larval stage (L4) of male development. The CEM neurons start to express sensory function genes upon sexual maturation at the L4 stage (Wang 2010). To avoid protein accumulation as secondary consequence of aging, we examine PKD-2 and CIL-7 localization at L4 stage to directly detect ciliary localization or EV shedding defects. In wild-type L4 cilia, PKD-2 and CIL-7 reached the ciliary tip and were shed into the environment (Fig. 3a). In *klp-6* mutants, both PKD-2 and CIL-7 entered cilia, but failed to locate distally and were not shed in environmental EVs (Fig. 3 b, d, e). In *klp-6* mutant cilia, PKD-2 formed small protrusions along the areas enriched with CIL-7 (Fig. 3b), supporting the hypothesis that enrichment of the CIL-7 protein drives PKD-2 EV biogenesis. The absence of strict colocalization between PKD-2 and CIL-7 suggested different ciliary location mechanisms.

**Figure 3.**
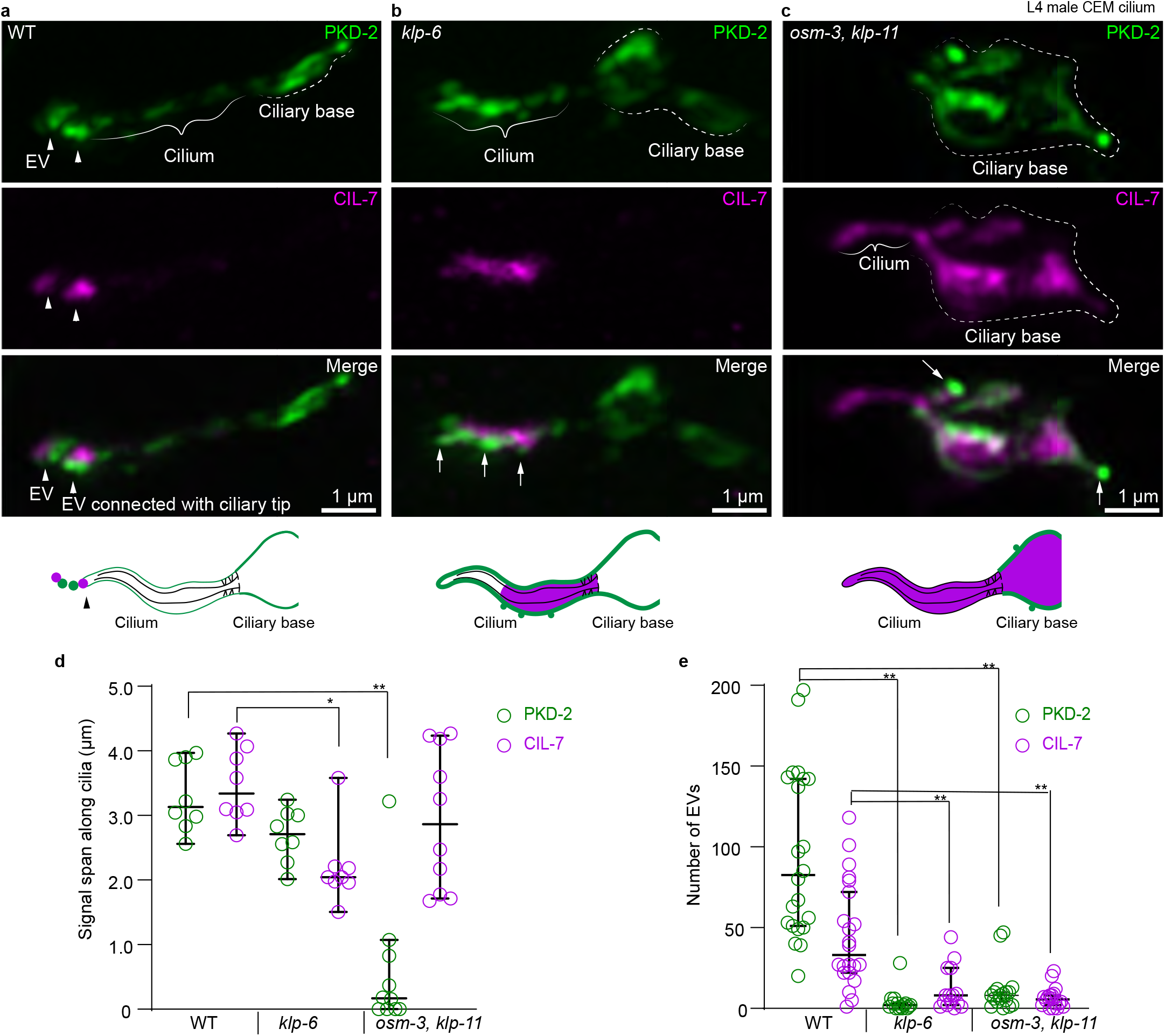
PKD-2 and CIL-7 require kinesin motors for their localization at the ciliary tip. **a-c**, Representative images of L4 male CEM cilia of wild type (a), kinesin-3 *klp-6* mutant (b), and kinesin-2 *osm-3 klp-11* double mutant (c). Top to bottom: PKD-2, CIL-7, merged channel, and cartoon summarizing the observed PKD-2 and CIL-7 localization. **a**, In the wild type, PKD-2::GFP puncta are observed at the ciliary membrane and are enriched at the ciliary tip. The CIL-7 is mostly enriched at the ciliary tip at places of active EV biogenesis. **b**, In the kinesin-3 *klp-6* mutant, PKD-2 and CIL-7 fail to reach the ciliary tip and remain ectopically enriched at the ciliary shaft. Note that PKD-2::GFP is organized into small protrusions adjacent to CIL-7 enriched areas. **c**, In the kinesin-2 *osm-3 klp-11* double mutant, PKD-2 fails to enter <u>stunted cilia</u> and accumulates considerably at the ciliary base. In contrast, CIL-7 locates to the cilium and abnormally accumulates at the ciliary base. The CIL-7 enrichment at the base corresponds to the formation of PKD-2 enriched protrusions, similar to phenotype observed at the ciliary shaft of the *klp-6* mutant (**b**). **d**, Quantification of the length of PKD-2 and CIL-7 along the cilia of wild type, *klp-6*, and *osm-3 klp-11* animals. CIL-7, but not PKD-2, traverses a significantly shorter distance in the *klp-6* mutant compared to wild type. PKD-2 localization is significantly affected in the *osm-3 klp-11* double mutant, but not in the *klp-6* mutant. **e**, Quantification of number of EVs labeled by PKD-2 and CIL-7 in late L4 male tail molting cuticle. Each data point represents total EV numbers in one animal. *klp-6* and *osm-3 klp-11* mutants are defective in both PKD-2 and CIL-7 EV release. Median values with 95% confidence intervals are indicated. \**p* < *0.05*, ** *p*<*0.01 by* Kruskal-Wallis test with Dunn’s multiple comparison, n=8, 8, and 10 for wild type, *klp-6,* and *osm-3 klp-11* in **d**, n=22, 15 and 18 for wild type, *klp-6* mutant, and *osm-3;klp-11* double mutant in **e**. Scale bars are 1 μm.

To determine how PKD-2 and CIL-7 enter cilia, we imaged their fluorescent reporters in the mutants with disruption in the intraflagellar transport (IFT) motors, homodimeric kinesin-2 OSM-3 (KIF17 in mammals) and heterotrimeric kinesin-2 KLP-11 (Fig. 3 c). In the *osm-3 klp-11* mutant, CIL-7 but not PKD-2 was able to enter the cilium, suggesting that PKD-2 ciliary entry was IFT-dependent whereas CIL-7 ciliary entry was not. *osm-3 klp-11* mutants displayed substantial CIL-7::tagRFP accumulation at the ciliary bases that correlated with PKD-2::GFP membrane protrusions. This phenotype resembled the PKD-2::GFP protrusions in CIL-7-enriched areas in *klp-6* mutant cilia (Fig. 3b). The *osm-3 klp-11* mutant did not produce environmental PKD-2 and CIL-7 EVs (Fig. 3e). Thus, PKD-2 EV biogenesis at the ciliary tip required functional IFT kinesin motors to enter cilia. Once in the cilium, PKD-2 employed CIL-7- and KLP-6-dependent mechanisms to gain enrichment at the ciliary tip and to be shed into environmental EVs.

What is the physiological significance of the independent sorting of PKD-2 and CIL-7 into environmental EVs? A known stimulant for male-specific sensory neurons is presence of mating partners. Additionally, PKD-2-carrying EVs are directly transferred from male sensory cilia to the hermaphrodite vulva during mating. We therefore tested whether exposure to hermaphrodites influenced abundance of PKD-2 and CIL-7 EVs produced by males (Fig. 4a). Upon male exposure to hermaphrodites, the ratio of PKD-2 EV to CIL-7 EV shedding increased compared to isolated, virgin males (Fig. 4b). These results indicate that cilia modulate EV cargo composition in response to sensory stimulation.

**Figure 4.**
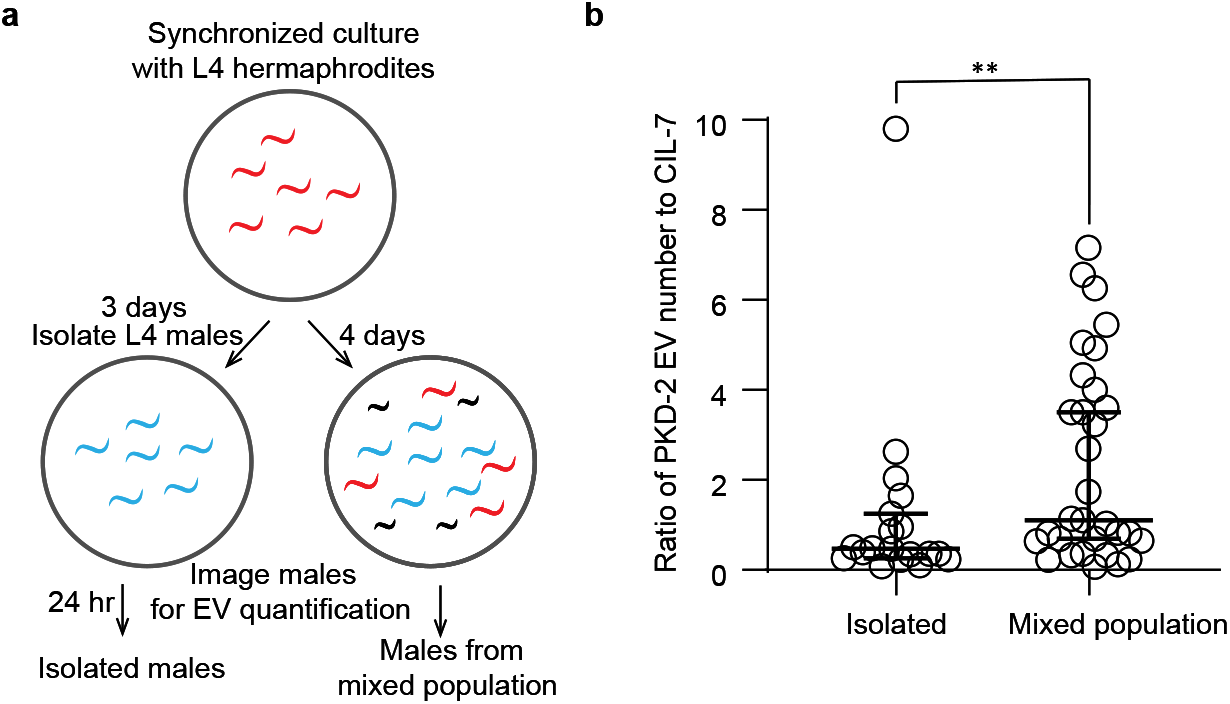
Ciliary tip EV biogenesis is altered by environmental cues. **a**, Scheme of experimental design for PKD-2 and CIL-7 EVs quantification in the presence and absence of mating partners. Males were isolated from a mixed population at the larval stage L4 and were imaged as adult virgin males alongside adult males picked from the mixed population with adult hermaphrodites. **b**, The ratio of PKD-2 to CIL-7 EV numbers is significantly increased in males cultured in mixed population as compared to isolated virgin males. **P<0.01 by Mann Whitney test, n=19 for isolated and 31 for mixed population conditions.

## Discussion

Cilia produce EVs from base and tip. We used Airyscan super-resolution microscopy to visualize and monitor ciliary EV shedding in vivo, which permitted study of EVs at an unprecedented spatiotemporal level. We show that PKD-2 EVs are shed at two distinct locations – the ciliary tip and the periciliary membrane. PKD-2 tip shedding from the ciliary tip is facilitated by the myristoylated coiled-coil CIL-7 protein that becomes enriched along the sites of PKD-2 EV biogenesis. Our study positions ciliary trafficking as a lever for controlling polycystin-2 ciliary tip EV shedding that requires IFT kinesin-2 motors and cell-specific kinesin-3 KLP-6.

The periciliary membrane acts as a novel compartment for ciliary base EV shedding. The antagonistic relationship between PKD-2 EV shedding from the ciliary tip versus the base is in accordance with our previous work showing that genetic disruption of PKD-2 environmental EV release correlates with massive PKD-2 ciliary base accumulation ^14,19,20,23^. The periciliary membrane is specialized region that regulates trafficking in and out of the cilium. Malfunction of IFT causes ciliopathies^24^. To date, the contribution of disrupted ciliary EV biogenesis into pathophysiology of ciliopathies remains unknown. Our study provides the first link between these two entities.

*C. elegans* EV-releasing cilia are specialized structurally and functionally. The cilia of *C. elegans* EV-releasing neurons (EVNs) protrude to the environment from cuticular pores ^25–27^ and shed EVs in response to mechanical pressure^28^. EVN ciliary membrane homeostasis of PKD-2 is modulated by a phosphoinositide 5-phosphatase^29^ and caveolin^30^. The tubulin code is required for the unique EVN ciliary ultrastructure and EV environmental release^19,23,31^, while kinesin-3 KLP-6 regulates intraflagellar transport and EV biogenesis at both the cilia base and tip^14,22^. The kinesin-3 motor family is characterized by their N-terminal motor domain, a forkhead-associated domain, and a tail possessing lipid-binding domains, the latter endows membrane-associated abilities^32^. The KLP-6 mammalian homolog KIF13B - via its lipid-binding C2 domain - maintains a caveolin-enriched membrane domain at the ciliary base in mammalian cells^33^. In addition to C2 domains, the KLP-6 tail contains positively charged amino acids^21^ that may enhance protein binding to negatively charged phospholipids in the ciliary membrane or polyglutamylated tubulin in ciliary axoneme^23,31^. We propose a mechanism whereby KLP-6 connects to both the ciliary membrane and axoneme and directs the myristoylated coiled-coil CIL-7 to distal ciliary tip for ciliary EV biogenesis. Coiled-coil proteins similar to CIL-7 mediate membrane fission by inserting the amphipathic helix into membrane bilayer^34^.

*C. elegans* ciliary tip EV shedding appears different than ciliary ectocytosis^9^ or ciliary tip decapitation in mammalian cell culture systems^10^. PKD-2-carrying EVs are released from the ciliary tip in one burst and range in number from several to hundreds (Extended Data Fig. 1a). By contrast, ciliary ectocytosis or decapitation events occur one at a time and over the course of several minutes. In IMCD3 cell culture, the KLP-6 homolog KIF13B is not required for ciliary ectocytosis of the neuropeptide Y receptor NPY2R^9^. The *Chlamydomonas* genome lacks kinesin-3 motors^35^, suggesting that different mechanisms drive ciliary tip shedding in these organisms.

Polycystin-1 and polycystin-2 localize to cilia and EVs throughout evolution. Our work provides a foundation for generation of testable hypotheses regarding polycystin EV biogenesis and function in human health and disease. Here, we show ciliary localization of PKD-2 positively correlates with its release in environmental EVs. Ciliary localization of polycystin-2 is essential for preventing cystogenesis in a mouse model^18^ and loss of cilia prevents cystogenesis in a Pkd2 mutant mouse^36^. We propose that ability of a cilium to package and shed polycystin-2 in EVs may be an important physiological process in the kidney and that defects in this process may contribute to cystogenesis.

*C. elegans* ciliary tip EV shedding is signal-dependent. Our data indicate that cilia fine-tune ciliary EV signaling for distant and local animal-to-animal communication. Adjustment of ciliary EV cargo ratios in response to sensory stimuli implies that neurons use sensory cilia to control outbound EV signaling. Heretofore cilium has been considered an organelle with two functions: sensation and/or motility. Our work provides a foundation for the third conserved function of the cilium – EV shedding for distant intercellular communication in multicellular organisms.

## Materials and Methods

No statistical methods were used to predetermine sample size. The experiments were not randomized. The investigators were not blinded to allocation during experiments and outcome assessment.

### Nematode culture and strains

*C. elegans* culture and genetics were performed as described (Brenner 1974). *C. elegans* strains were maintained at 22°C on nematode growth medium (NGM) plates seeded with *Escherichia coli* (OP50 strain) as a food source. CIL-7::tagRFP transgene was constructed by Gibson Cloning with forward primer 5’-ATG GAT ACG CTA ACA ACT TGG GCT GGG AGT CGA TAC ATG GT-3’ and reverse primer 5’-CAG CTC TTC GCC CTT AGA CAC ATG ATG TGC AGA CTT CTT CTT TC-3’ used to amplify it from genomic DNA of the natural isolate N2 as a template. Amplified fragments were cloned to the pPD95.75 vector backbone (gift from Andrew Fire, Addgene plasmid #1494) modified to carry tagRFP. Generation of extra-chromosomal array transgenes was carried out using standard procedures. The procedure included injection of a plasmid cocktail into the *pha-1(e2123ts)III; him-5 V* strain. The plasmid cocktail contained the transgene-of-interest at 10 ng/μL and a rescue transgene for the temperature-sensitive mutation in the *pha-1* gene (aka pBX1 *pha-1*+) at 100 ng/μL. Obtained stable line *myEx888[CIL-7::tagRFP*+*pBX1]* was further crossed to the *klp-6(my8)III* mutant to generate the PT3383 strain, and to the *osm-3(p802) klp-11(tm324)IV* double mutant to generate the PT3380 strain.

### Airyscan super-resolution microscopy

Super-resolution imaging was performed on the Zeiss LSM880 confocal system equipped with Airyscan Super-Resolution Detector with Fast Module, 7 Single Photon Lasers (405, 458.488, 514, 561, 594, 633nm), Axio Observer 7 Motorized Inverted Microscope, Motorized X-Y Stage with Z-Piezo, T-PMT. Planes of captured Z-stacks were distanced at 0.09 to 0.185 μm for optimal 3D rendering. Image analysis and processing was performed using Zen Blue 2.0 and Zen Black 2.0. The Imaris 9.5 (Bitplane, US) package was used to generate surface rendered three-dimensional reconstructions from the Airyscan processed images.

### Mounting males for super resolution live animal imaging

Males were anesthetized with 10 mM levamisole solution in the M9 buffer. To completely immobilize the male, we used thin agarose pads, prepared from 10% melted agarose (Sigma #A9539) and Millipore water. Melting of 10% agarose was achieved by pulse heating in a microwave in 3 second intervals for a total of 1 minute or until dry agarose pieces were no longer visible. Once melted, the 10% agarose was incubated at 95°C for about 10 minutes to allow air bubbles to escape. The melted agarose was dispensed onto glass slides in 100 μL droplets and the droplets were immediately pressed with a second glass slide to make a thin gel pad. The agarose pads were made in batches and stored in a sealed glass slide container at room temperature for up to a week. Prior to use, one glass slide was removed, the agarose pad was dried at room temperature until excess water was no longer visible, yet pad remained moist. Once achieved, the agarose pad was cut into 1 cm x 1 cm square to remove dried edges and ensure even surface. Males were placed in 1 μL droplet of 10 mM Levamisole dispensed on a 18 × 18 cm coverslip, then the coverslip with males was flipped on the center of the agarose pad. Semi-dry agarose pads held the males tightly under coverslip by capillary action and minimized worm moving.

### Late L4 and young adult male preparation for super-resolution imaging the CEM cilia

About 100 - 200 L4 males were picked from a healthy *C. elegans* culture to a new seeded plate prior to imaging. Three hours later young adult males with completely developed tail fan and rays were selected for imaging. Late L4 males were picked just prior to molting when cuticle was covering the almost fully mature tail. Young adult males were defined by the absence of the molt cuticle covering their tails.

### PKD-2 and CIL-7 EV number and ratio quantification for CEM cilia

Populations were synchronized by picking 10 L4 hermaphrodites on new seeded NGM plates. Three days later some L4 males were isolated from the mixed population to deprive males of exposure to their mating partners during final steps of their sexual maturation. The isolated virgin males were imaged the next day (24 hours later) to quantify their capacity to release EVs. This imaging was done in parallel with imaging of adult males that were exposed to hermaphrodites until imaging time. Imaging was performed in 1 mM Levamisole prepared with ddH_2_O on 5% agarose pads. Colocalization of PKD-2 and CIL-7 was defined by a distance between centroids of the fluorescent areas being less than 200 nm using a custom python script. The script is available upon request.

### Statistical analysis

The Prism software package (GraphPad Software 8) was used to carry out statistical analyses. Information about statistical tests, p values and n numbers are provided in the respective figures and figure legends.

### Reporting summary

Further information on research design is available in the Nature Research Reporting Summary linked to this paper.

### Data availability

All relevant data are available and/or included with the manuscript as Source Data or Supplementary Information.

---

**Supplemental Table 1.** Strains used in this study.

| Strain name | Genotype | Reference |
| --- | --- | --- |
| PT1435 | *pha-1(e2123) III; myIs1 IV; him-5(e1490) V; myEx552 [Ppkd-2::TBB-4::tdTomato]* | Jauregui 2008 |
| PT823 | *pha-1(e2123) III; myIs1 [PKD-2::GFP+Punc-122::GFP]IV; him-5(e1490)V; myEx431 [Pnphp-1::NPHP-1::dsRED]* | Jauregui 2008 |
| PT621 | *him-5(e1490) myIs4 [PKD-2::GFP+Punc-122::GFP]V* | Bae 2006 |
| PT3112 | *pha-1(e2123) III; him-5(e1490) V myIs4V; myEx888[CIL-7::tagRFP + pBX1].* | This work |
| PT3383 | *klp-6(my8) III; him-5(e1490) myIs4V; myEx888[CIL-7::tagRFP + pBX1].* | This work |
| PT3380 | *pha-1(e2123) III; osm-3(p802) klp-11(tm324)IV; him-5(e1490) myIs4V; myEx888[CIL-7::tagRFP + pBX1].* | This work |
| PT2681 | *cil-7(tm5848) I; myIs1[PKD-2::GFP+Punc-122::GFP] IV; him-5(e1490)V* | Maguire 2015 |

**Extended Data Figure 1.**
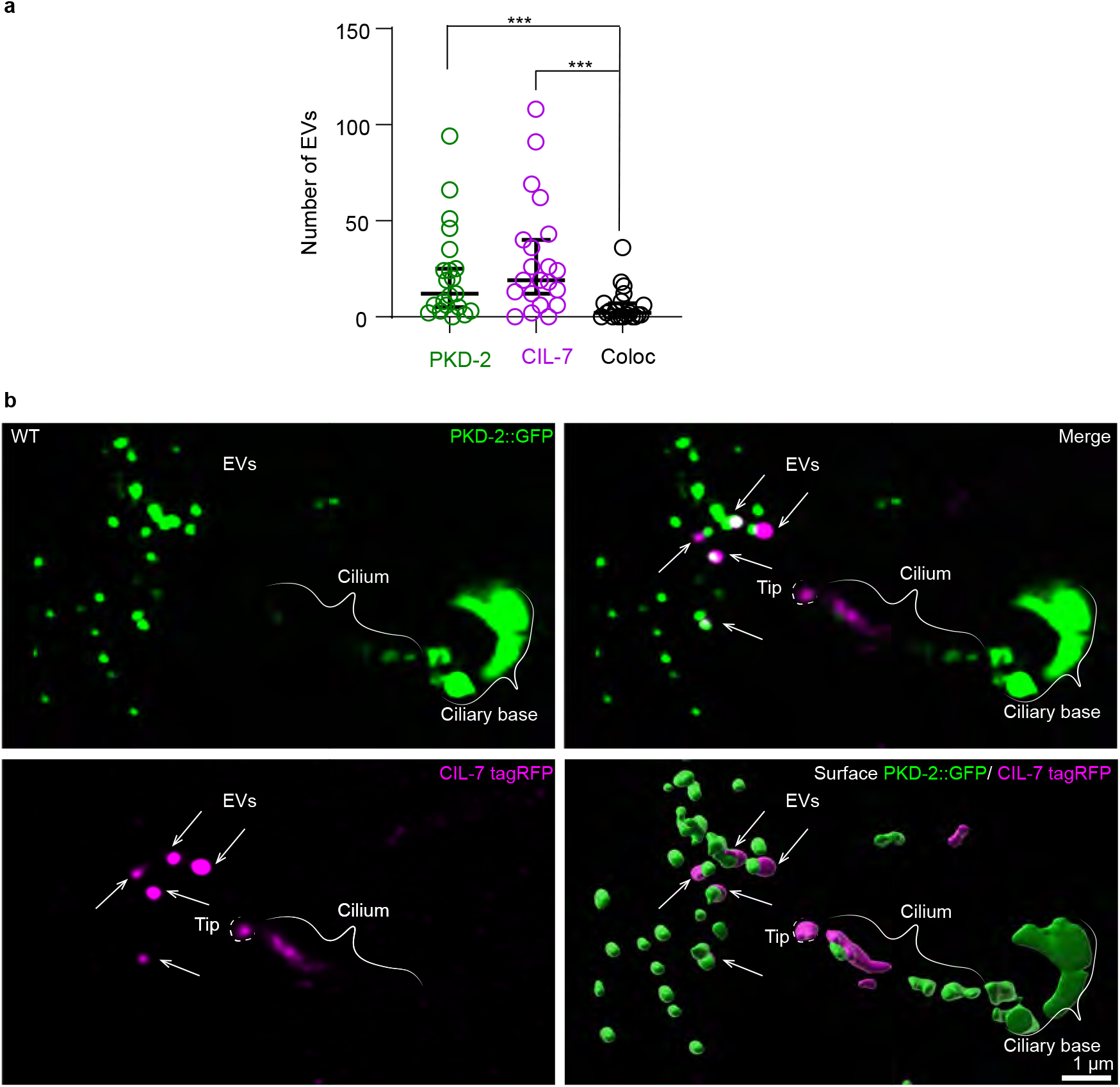
PKD-2 and CIL-7 are sorted to distinct environmental EVs. **a**, Analysis of colocalization of PKD-2 and CIL-7 on environmental EVs released from ciliary tips of adult wild-type isolated males. PKD-2 and CIL-7 are mostly sorted into separate EVs. The graph contains paired data where each data point represents number of EVs released from a single male head. Median values with 95% confidence intervals are indicated. *** p<0.001 by Friedman test, n=21. **b**, Environmental EVs are observed in the vicinity of the releasing wild-type cilium. The imaged is captured at the moment of CIL-7 enrichment at the ciliary tip (outlined by dashed white line) that is about to separate from the cilium. White arrows point to CIL-7 EVs. Note each of the CIL-7 EVs stays connected with PKD-2 EVs after their release. Scale bar 1 μm.

**Extended Data Figure 2.**
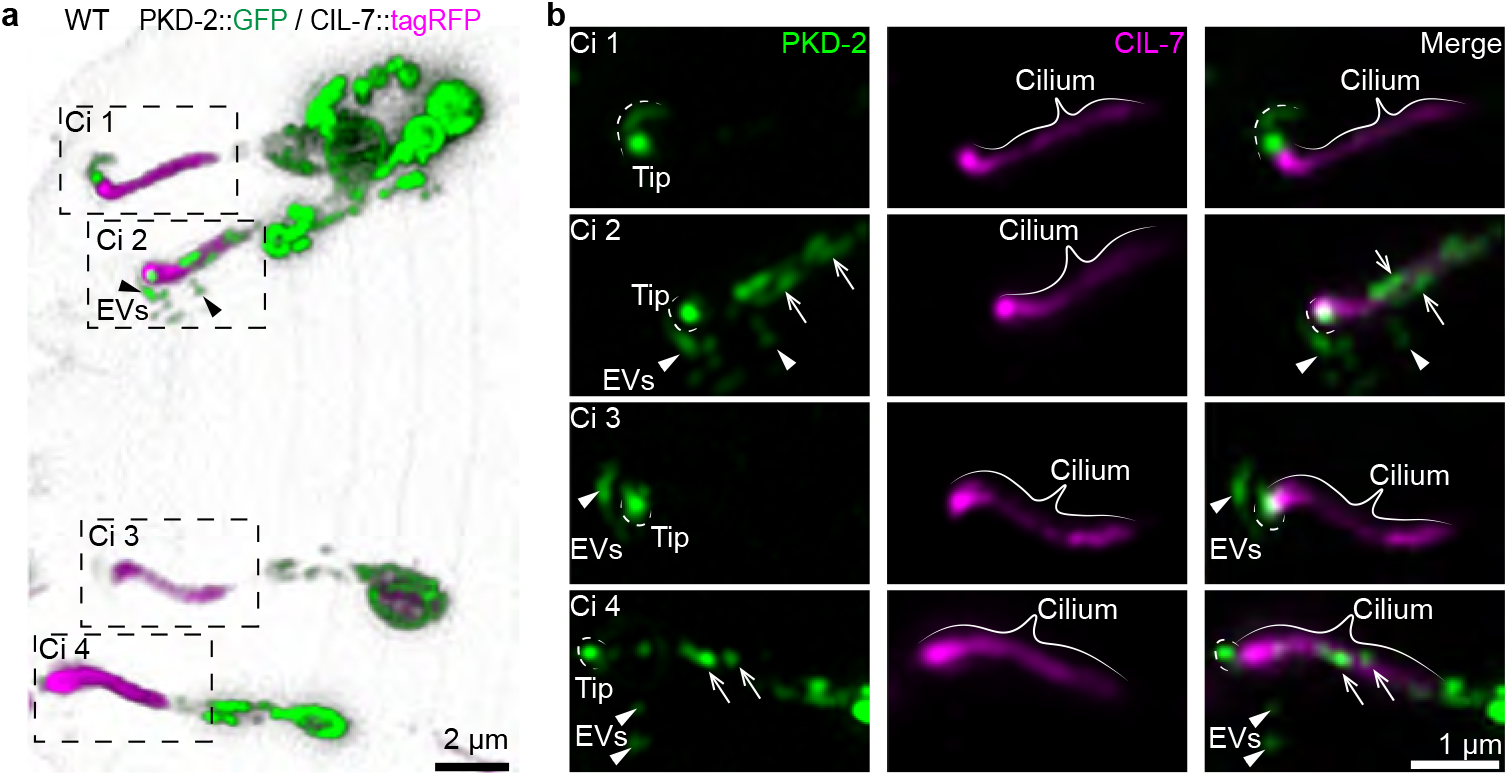
CIL-7-mediated enrichment of PKD-2 at the ciliary tip is required for biogenesis of environmental EVs. **a**, Three-dimensional rendering model of a male head with all four CEM cilia showing PKD-2::GFP and CIL-7::tagRFP distributions. Each cilium is numbered and shown on close-up images on panel **b**. **b**, PKD-2::GFP enrichment at the ciliary tip is outlined by white dashed lines. White arrowheads indicate environmental EVs. White arrows indicate PKD-2::GFP captured along the ciliary shaft of Ci 2 and Ci 4. Scale bar 2 μm for a, 1 μm for b.

**Extended Data Fig. 3.**
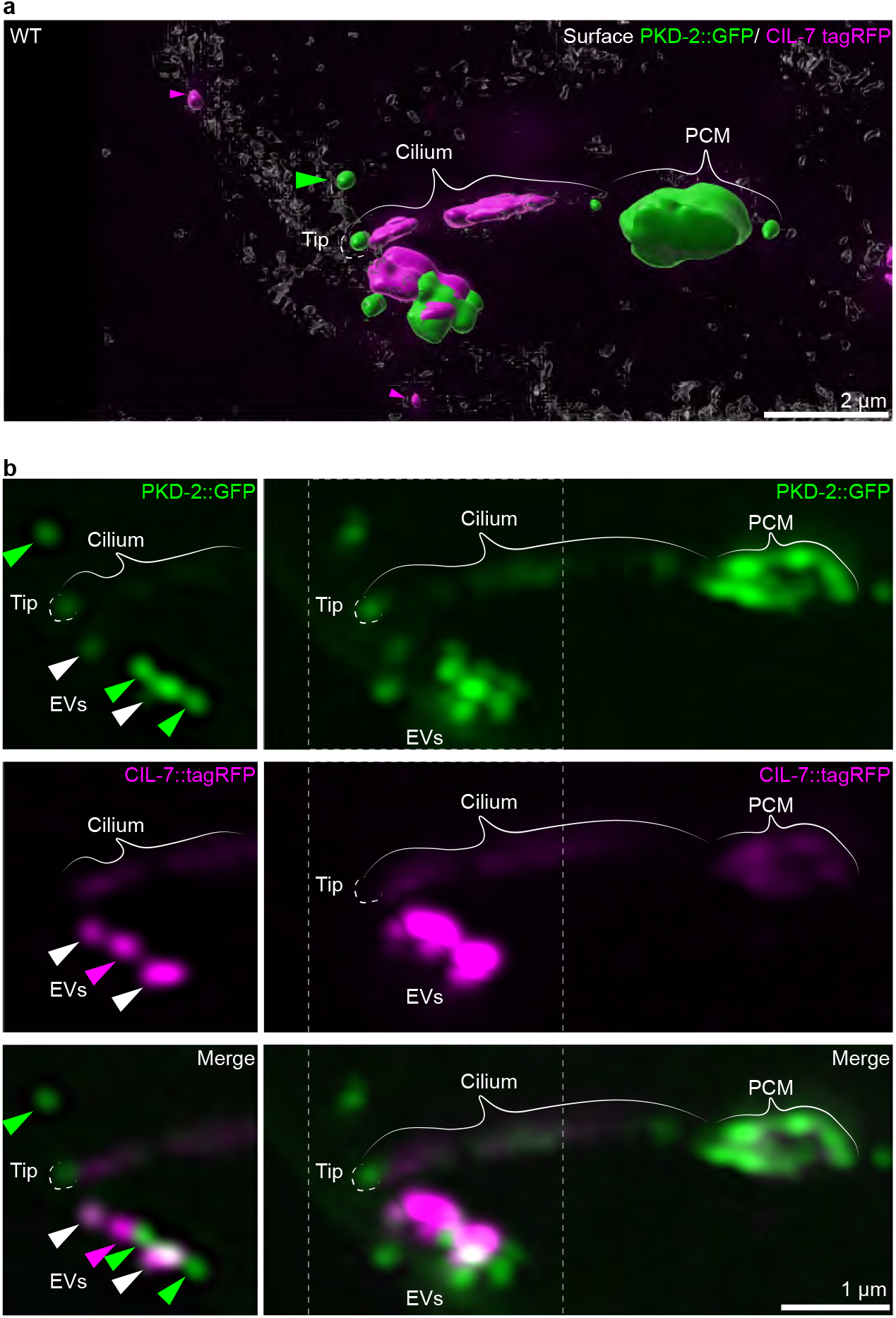
Simultaneous release of PKD-2 and CIL-7 environmental EVs. **a**, Surface rendered image of PKD-2::GFP and CIL-7::tagRFP distribution along the cilium of a young adult male. PKD-2::GFP is enriched at the ciliary base and the ciliary tip. CIL-7::tagRFP is captured at a moment of forming a larger protrusion from the ciliary tip that still houses multiple PKD-2::GFP EVs not completely separated yet from the CIL-7-enriched launchpad. **b**, Split view of channels used to produce the surface rendered image on panel **a**. Left column shows a select focal plane outlined with white dashed lines on the images positioned in the right column, which contains orthogonal projections of the entire Z-stack. On the single focal plane PKD-2 and CIL-7 EVs are alternating with each other within one string budding from the ciliary tip. Green arrowhead indicates the sole PKD-2 environmental EV; magenta arrowheads indicate the sole CIL-7 environmental EVs; white arrowheads show points of PKD-2 and CIL-7 colocalization. Scale bar, 2 μm in a, 1 μm in b.

